# Piezo1 channel agonist mimics high glucose as a stimulator of insulin release

**DOI:** 10.1101/455832

**Authors:** Vijayalakshmi Deivasikamani, Savitha Dhayalan, Romana Mughal, Asjad Visnagri, Kevin Cuthbertson, Jason L Scragg, Tim S Munsey, Hema Viswambharan, Richard Foster, Asipu Sivaprasadarao, Mark T Kearney, David J Beech, Piruthivi Sukumar

## Abstract

**Objective:** Glucose and hypotonicity induced cell swelling stimulate insulin release from pancreatic β-cells but the mechanisms are poorly understood. Recently, Piezo1 was identified as a mechanically-activated nonselective Ca^2+^ permeable cationic channel in a range of mammalian cells. As cell swelling induced insulin release could be through stimulation of Ca^2+^ permeable stretch activated channels, we hypothesised a role for Piezo1 in cell swelling induced insulin release.

**Methods:** Two rat β-cell lines (INS-1 and BRIN-BD11) and freshly-isolated mouse pancreatic islets were studied. Intracellular Ca^2+^ measurements were performed using the fura-2 Ca^2+^ indicator dye. Piezo1 agonist Yoda1, a competitive antagonist of Yoda1 (Dooku1) and an inactive analogue of Yoda1 (2e) were used as chemical probes. Piezo1 mRNA and insulin secretion were measured by RT-PCR and ELISA respectively.

**Results:** Piezo1 mRNA was detected in both β-cell lines and mouse islets. Yoda1 evoked Ca^2+^ entry which was inhibited by Yoda1 antagonist Dooku1 as well as other Piezo1 inhibitors gadolinium and ruthenium red, and not mimicked by 2e. Yoda1, but not 2e, stimulated Dooku1-sensitive insulin release from β-cells and pancreatic islets. Hypotonicity and high glucose increased intracellular Ca^2+^ and enhanced Yoda1 Ca^2+^ influx responses. Pre-treatment with ruthenium red significantly reduced hypotonicity induced insulin release from β-cells and pancreatic islets.

**Conclusion:** The data show that Piezo1 channel agonist induces insulin release from β-cell lines and mouse pancreatic islets suggesting a role for Piezo1 in cell swelling induced insulin release. Hence Piezo1 agonists have a potential to be used as enhancers of insulin release.

## 1 Introduction

Secretion of insulin by pancreatic β-cells is vital for maintaining blood glucose and energy homeostasis. Although insulin biosynthesis is controlled by multiple factors, glucose metabolism is the most important physiological event that stimulates insulin secretion [1]. Physiologically, glucose enters the β-cells through an insulin independent process (involving glucose transporter 1, GLUT-1) where it undergoes glycolysis, enters the TCA cycle and results in the generation of ATP. An increased ATP/ADP ratio leads to closure of the ATP-dependent potassium (K_ATP_) channels which induces membrane depolarisation which in turn, activates voltage-dependent Ca^2+^ channels (VDCC), leading to influx of Ca^2+^. Elevated intracellular Ca^2+^ forms the trigger for insulin release. Insulin is then released into the circulation by the fusion of granules with the cell membrane and exocytosis [2]. Apart from the conventional ATP-K_ATP_-VDCC pathway of insulin release in response to glucose stimulation, osmotic cell swelling resulting from increased glucose levels or hypotonicity has also been reported to induce insulin secretion [3]–[6]. Glucose induced cell swelling is suggested to be because of lactate accumulation and/or activation of Na^+^/H^+^ and Cl^−^/HCO_3_ exchangers [7]. Such insulin secretion was not completely inhibited by K_ATP_ blockers [7], [8]. Cell swelling has been suggested to stimulate insulin release by activating anionic channels in pancreatic β-cells namely the volume sensitive chloride (Cl^-^) channels, thus depolarizing the cell membrane and increasing the cytosolic Ca^2+^ concentration through the activation of VDCC [3], [4], [9], [10]. However, some investigators have argued against this as hypotonically induced insulin secretion persisted even in the presence of Cl^-^ channel blockers such as niflumic acid and DIDS [6], [11]. Thus it is possible that a mechanism independent of Cl^-^ channels is involved in hypotonically induced insulin secretion. Moreover, a study on rat pancreatic islets *in vitro* has revealed that glucose and hypo-osmotic cell swelling induce insulin secretion through distinct signal transduction pathways [12]. Osmotic cell swelling mechanically stretches the plasma membrane and hence the role of stretch activated cation channels was investigated [13]. The latter study revealed that hypotonicity induced osmotic cell swelling in rat pancreatic β-cells results in the activation of certain stretch activated cationic channels which results in Ca^2+^ entry and insulin release. However, no report currently exists on the identity of stretch activated channels in pancreatic β-cells [7].

Recently, a novel type of stretch activated Ca^2+^ permeable non selective cationic channels named Piezo1 has been identified in various mammalian cells [14]. Piezo1 is expressed in the lungs, bladder, kidney, endothelial cells, erythrocytes, skin, periodontal ligament cells and elsewhere [15]. Piezo1 channels are responsible for mechanically activated cationic currents in numerous eukaryotic cell types, converting mechanical forces to biological signals [16]. Piezo1 has been reported to play a significant role in endothelial cell biology, collecting duct osmoregulation, urothelial pressure sensing, cell migration and erythrocyte volume regulation [17]–[21]. Piezo1 is a large integral membrane protein with 38 transmembrane segments which assemble as trimers to form ion channels [22], [23]. Piezo1 can be specifically activated by Yoda1, a key compound used to study Piezo1 regulation and function, and is inhibited by common non-specific small molecule inhibitors such as ruthenium red (RR) and gadolinium ions [24], [25]. Recently, a Yoda1 analogue named Dooku1 has been identified which antagonizes Yoda1 activation of Piezo1 [26].

We hypothesised that Piezo1 could contribute to stretch induced Ca^2+^ influx induced by cell swelling in β-cells, and thereby regulate insulin secretion. In order to test our hypothesis, two rat β-cell lines (INS-1 and BRIN-BD11) were used as model β-cell line and mouse pancreatic islets were also used in conjunction with chemical modulators of Piezo1. Our results indicate that Piezo1 channels have a significant role in hypotonicity/cell swelling induced insulin release from β-cells.

## 2 Materials and Methods

### 2.1 Chemicals and Solutions

Unless stated otherwise, all commercially available chemicals were obtained from Sigma-Aldrich. Stocks of chemicals were reconstituted in DMSO and stored at −20°C unless stated otherwise. Fura-2-AM (Molecular Probes) was dissolved at 1 mM concentration. Pluronic acid F-127 was stored at room temperature at 10% w.v**^-^**^1^ in DMSO. Yoda1 (Tocris) was stored at 10 mM. Yoda1 antagonist (Dooku1) and non-functional Yoda1 analogue (2e) were synthesized and purified and prepared as 10 mM stock solutions [26]. Standard bath solution (SBS) contained (in mM): 140 NaCl, 5 KCl, 1.2 MgCl_2_, 1.5 CaCl_2_, 8 glucose and 10 HEPES titrated to pH 7.4 using NaOH. For low/high glucose experiments, SBS contained (in mM): 85 NaCl, 5 KCl, 1.2 MgCl_2_, 1.5 CaCl_2_, 2.8/17.8 glucose, 10 HEPES and 100/85 D-mannitol titrated to pH 7.4 using NaOH. For hypo/normotonic experiments, SBS contained (in mM): 85 NaCl, 5 KCl, 1.2 MgCl_2_, 1.5 CaCl_2_, 2.8 glucose, 10 HEPES and 0/100 Mannitol titrated to pH 7.4 using NaOH. Krebs-Ringer bicarbonate HEPES (KRBH) buffer contained (in mM): 120 NaCl, 1.25 KH_2_PO_4_, 1.25 MgSO_4_, 2.68 CaCl_2_, 5.26 NaHCO_3_ and 10 HEPES. Zero-, low- and high-glucose KRBH contained 0, 2.8, 17.8 mM glucose. Osmolality was maintained across all three buffers by adding an equivalent amounts of D-mannitol to zero and low glucose KRH.

### 2.2 Cell culture

INS-1 832/13 (referred as INS-1) cells derived from rat insulinoma were cultured in RPMI 1640-GlutaMAX-I (Gibco, USA) medium containing 10% heat-inactivated foetal calf serum, penicillin (100 U/ml), streptomycin (100 μg/ml), 1 mM sodium pyruvate, 50 μM 2-mercaptoethanol and 10 mM HEPES. BRIN-BD11 cells derived from rat pancreatic islets were grown in RPMI 1640 -GlutaMAX-I (Gibco, USA) medium containing 10% heat-inactivated foetal calf serum, penicillin (100 U/ml), streptomycin (100 μg/ml). Cultures were maintained at 37 °C under 5% CO_2_ and a humidified atmosphere. Subcultures were established once every 3-4 days by trypsin/EDTA treatment.

### 2.3 Mouse pancreatic islet isolation

Islets were isolated following the published protocol [27], [28]. Briefly, mice were sacrificed and the pancreata harvested in sterile HEPES buffer. In a sterile hood, the pancreas was washed twice with phosphate buffer saline (PBS) supplemented with 1% penicillin (100 U/ml), streptomycin (100 μg/ml). The pancreas was then minced thoroughly in a glass petri dish and digested using 0.1% of collagenase-IV (Sigma) prepared in serum-free RPMI1640 media. The digestion was carried out for 5-7 minutes at 37 °C under 5% CO_2_ with frequent shaking; 2 ml of fetal bovine serum was added to neutralize the collagenase. The digested tissue was then centrifuged at 800 rpm for 10 minutes. The resulting pellet was resuspended in DMEM medium containing 10% foetal calf serum, penicillin (100 U/ml), streptomycin (100 μg/ml) and seeded in a 60mm Petri dish and cultured for 48 hours at 37 °C under 5% CO_2_ and a humidified atmosphere. C57BL/6 mice aged 2-5 months were used to isolate pancreas, which were conducted in accordance with accepted standards of humane animal care under United Kingdom Home Office Project license No. P606320FB.

### 2.4 RT-PCR

Cells or islets were lysed using TRIzol reagent (Life Technologies), and total RNA was isolated according the manufacturer’s protocol. After DNaseI treatment, RNA was quantified spectrometrically and 1 µg of RNA was reverse transcribed using OligodT primers (Life Technologies) following the manufacturer’s protocol. PCR was performed using *Piezo1* specific intron spanning primers (mouse Piezo1: Forward-5’CTGGACCAGTTTCTGGGACAA3’ and reverse-5’AGCCTGGTGGTGTTAAAGATGT3’; rat Piezo1 Forward-5’TTCTTCGGGGTGGAGAGGTA3’ and reverse-5’TGTCACCATGTGGTTAAGGATG3’). PCR products were sequenced for confirmation of the products and resolved electrophoretically on 1.5% agarose gel for presentation purposes.

### 2.5 Intracellular Ca^2+^ measurements

INS-1 or BRIN-BD11 cells were plated in poly-D-lysine coated 96-well plates (Corning, NY, USA) to 80-90 % confluence 24 hr before experiments. Prior to recordings, cells were incubated for 1 hr at 37 °C in 2.5 µM fura-2AM dispersed in SBS. The cells were washed for 0.25 hr in SBS at room temperature. Measurements were made on a 96-well bench-top scanning fluorimeter (FlexStation III) with SoftMax Pro v5.4.5 (Molecular Devices, Sunnyvale, CA, USA). Intracellular Ca^2+^ was indicated as the ratio of fura-2 emission (510 nm) intensities for 340 and 380 nm excitation. Experiments were performed at room temperature (21±2 °C). For Piezo1 inhibition, hypotonicity and high glucose experiments, the inhibitors/modified bath solutions were added on to the cells at the wash point (after fura-2 loading) and the same concentrations were maintained throughout the experiment.

### 2.6 Insulin secretion stimulation and assessment

Cells or islets were washed with zero glucose KRBH buffer for 1 hr at 37 °C. Dooku1 pre-treatment was achieved by replacing the wash buffer with fresh buffer containing 10 µM Dooku1 during the last 15 min of the wash period. After washing, fresh low glucose KRH buffer containing test chemicals (Yoda1 and the analogues) was added onto the cells/islets and incubated for 1 hr at 37 °C. High glucose KRBH and hypotonic SBS were used for measuring high glucose and hypotonicity induced insulin release. Cells are exposed to circular shear stress by placing the culture plates on an orbital rotating platform (Grant Instruments) housed inside the incubator for 1 hour. The radius of orbit of the orbital shaker was 10 mm and the rotation rate was set to 210 rpm, which caused swirling of the buffer over the cell surface volume. The movement of fluid due to orbital motion represents a free surface flow at the liquid-air interface, of 12 dyne/cm^2^. Next, the supernatant was collected and stored at -80°C until analysis. RR pre-treatment was done by replacing the wash buffer with fresh buffer containing 30 µM RR during last 15 min of the wash period. For islets, buffer change and supernatant collection was performed by centrifuging the islets at 800 rpm for 10 min. Insulin levels were determined using an ELISA kit according to the manufacturer’s protocol (from Crystal Chem and RayBio for mouse and rat respectively). For islets, the insulin level was normalised to the total protein of each sample. Protein quantification was carried out by the BCA assay (Pierce) after lysing the islets in the protein lysis buffer containing (in mmol/L), 50 HEPES, 120 NaCl, 1 MgCl_2_, 1 CaCl_2_, 10 NaP_2_O_7_, 20 NaF, 1 EDTA, 10% glycerol, 1% NP40, 2 sodium orthovanadate, 0.5 µg/mL leupeptin, 0.2 PMSF, and 0.5 µg/mL aprotinin (Invitrogen Cell Extraction Buffer).

### 2.7 Data analysis and Statistics

All data were analysed using Origin 2016 software. Results are expressed as mean ± standard error of mean (SEM) of at least 3 independent repeats. Comparisons within groups were made using paired Students t-tests and between groups using unpaired Students t-test, as appropriate; p <0.05 was considered statistically significant.

## 3 Results

### 3.1 INS-1 cells express Piezo1 and respond to Piezo1 agonist

Piezo1 mRNA was detected in INS-1 cells by RT-PCR (Figure 1A). Anticipating that functional channels are also expressed, the activity of Piezo1 chemical agonist Yoda1 on Ca^2+^ influx was tested on INS-1 cells. In a dose dependent manner Yoda1 induced increases in intracellular Ca^2+^ with an estimated EC_50_ of 4 µM (Figure 1B&C). Poor solubility of Yoda1 above 10 µM prevented accurate determination of the EC_50_. To investigate if the Ca^2+^ increase depends on influx of Ca^2+^ from extracellular space, rather than release from intracellular stores, effect of Yoda1 was tested in the absence of extracellular Ca^2+^. When there was no Ca^2+^ outside, Yoda1 did not increase the intracellular Ca^2+^ signal, indicating that it induces Ca^2+^ influx rather than store release (Figure 1D&F). The small increase observed above baseline in the absence of extracellular Ca^2+^ is most likely an artefact of solution application. Most β-cell lines including INS-1 cells express VDCC which are essential for the classical insulin secretion pathway. Therefore cells were pre-treated with the VDCC inhibitor nicardipine (10 µM for 30 min) before the addition of Yoda1. Nicardipine did not affect the Yoda1 stimulation of INS-1 cells, suggesting that the Yoda1 induced Ca^2+^ influx was not through VDCC (Figure 1E&F). Though Yoda1 is considered to be specific for Piezo1 channels, the risk of non-specific effects must be considered. Thus we have made additional determinations. The non-specific inhibitors Gd^3+^ and ruthenium red (RR), both at 30 µM concentration with 30 min pre-treatment, inhibited Yoda1 induced Ca^2+^ influx (Figure 1G&H). Moreover, the recently developed Yoda1 analogue (Dooku-1), which antagonises Yoda1 activation of Piezo1, significantly reduced Yoda1 induced Ca^2+^ influx (Figure 1I&K) and an inactive Yoda1 analogue (2e) failed to mimic the effect of Yoda1 (Figure 1J&K). These expression and pharmacological data suggest that INS-1 cells contain functional Piezo1 channels.

**Figure 1.**
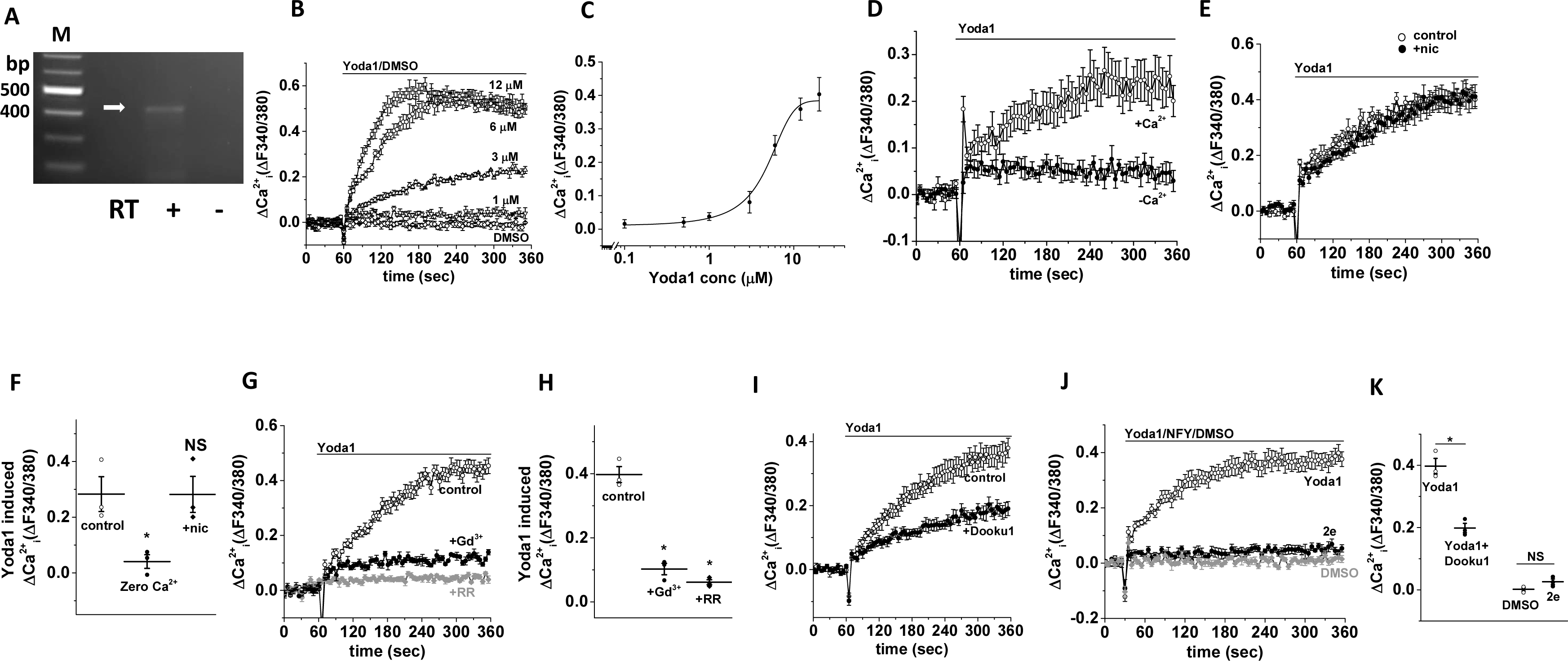
INS-1 cells express functional Piezo1 channels. **A)** Representative agarose gel electrophoresis picture with a single band at 473 bp (arrow) showing Piezo1 mRNA expression (RT: with (+) and without (-) reverse transcriptase; bp: base pairs; M: marker). **B)** Example data showing Yoda1 (increasing doses) or DMSO (control) induced Ca^2+^ influx in INS-1 cells. **C)** Mean data obtained from 5 independent experiments of the type shown in (**B**) showing the concentration-response curve for Yoda1 effect. **D&E)** Example data showing the effect of zero extracellular Ca^2+^ (**D**) and nicardipine (**E**) pre-treatment on Yoda1-mediated Ca^2+^ response in INS-1 cells. **F)** Mean data obtained from 3 independent experiments of the types shown in (**D&E**). **G)** Example data showing the effect of gadolinium (Gd^3+^) and ruthenium red (RR) pre-treatment on Yoda1-mediated Ca^2+^ response in INS-1 cells. **H)** Mean data obtained from 3 independent experiments of the types shown in (**G**). **I)** Example data showing the effect of Dooku1 pre-treatment on Yoda1-mediated Ca^2+^ response in INS-1 cells. **J)** Example data showing the effect of Yoda1, a non-functional Yoda1 analogue (2e) and DMSO (control) on Ca^2+^ influx in INS-1 cells. **K)** Mean data obtained from 3 independent experiments of the types shown in (**I&J**). Data are expressed as mean±sem; * denotes p<0.05 vs control; all data are representative of or mean obtained from 3 independent repeats.

### 3.2 Piezo1 agonist response in BRIN-BD11 cells

To further confirm the expression of functional Piezo1 channels in β-cell lines, we also investigated another β-cell line: BRIN-BD11 cells. These cells also expressed Piezo1 mRNA (Figure 2A). Similar to INS-1 cells,Yoda-1 induced Ca^2+^ influx occurred in BRIN-BD11 cells (estimated EC_50_ 9 μM; Figure 2B&C) and was inhibited by Gd^3+^ (30 μM) or Dooku1 (10 μM) (Figure 2D&E). The data suggest that BRIN-BD11 cells also contain functional Piezo1 channels.

**Figure 2.**
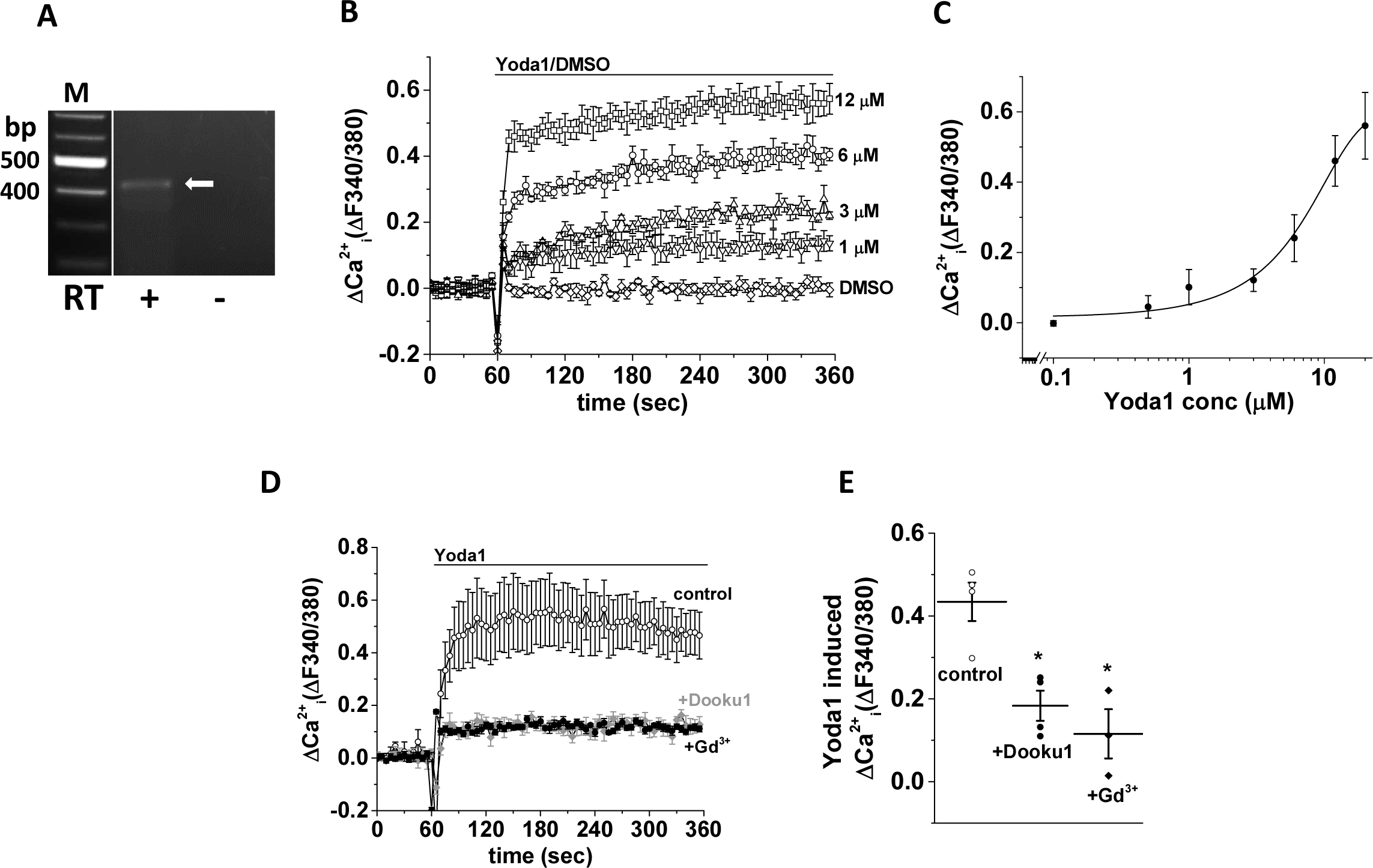
BRIN-BD11 cells express functional Piezo1 channels. **A)** Representative agarose gel electrophoresis picture with a single band at 473 bp (arrow) showing Piezo1 mRNA expression (RT: with (+) and without (-) reverse transcriptase; bp: base pairs; M: marker). **B)** Example data showing Yoda1 (increasing doses) or DMSO (control) induced Ca^2+^ influx in BRIN-BD11 cells. **C)** Mean data obtained from 3 independent experiments of the type shown in (**B**) showing the concentration-response curve for Yoda1 effect. **D)** Example data showing the effect of gadolinium (Gd^3+^) and Dooku1 pre-treatment on Yoda1-mediated Ca^2+^ response in INS-1 cells. **E)** Mean data obtained from 3 independent experiments of the types shown in (**D**). Data are expressed as mean±sem; * denotes p<0.05 vs control; all data are representative of or mean obtained from 3 independent repeats.

### 3.3 Stimulation of insulin release by osmotic or shear stress

An expectation of cells expressing Piezo1 is that they should be sensitive to osmotic stress caused by hypotonic solution and shear stress caused by fluid flow. Hypotonic solution is expected to cause cell swelling because of increased water entry into cells and specifically in β-cells, cell swelling leads to insulin release. Consistent with the presence of functional Piezo1 channels, hypotonicity increased basal and Yoda1-induced intracellular Ca^2+^ levels (Figure 3A-D). Furthermore hypotonicity caused insulin secretion which was suppressed by the non-specific Piezo1 channel inhibitor RR (Figure 3E). Exposing INS-1 cells to the physical force of shear stress (12 dyne/cm^2^ for 1 hour) also induced insulin release and this too was inhibited by RR (Figure 3F). The data provide further evidence of functional Piezo1 channels in β-cells and suggest a link to insulin secretion.

**Figure 3.**
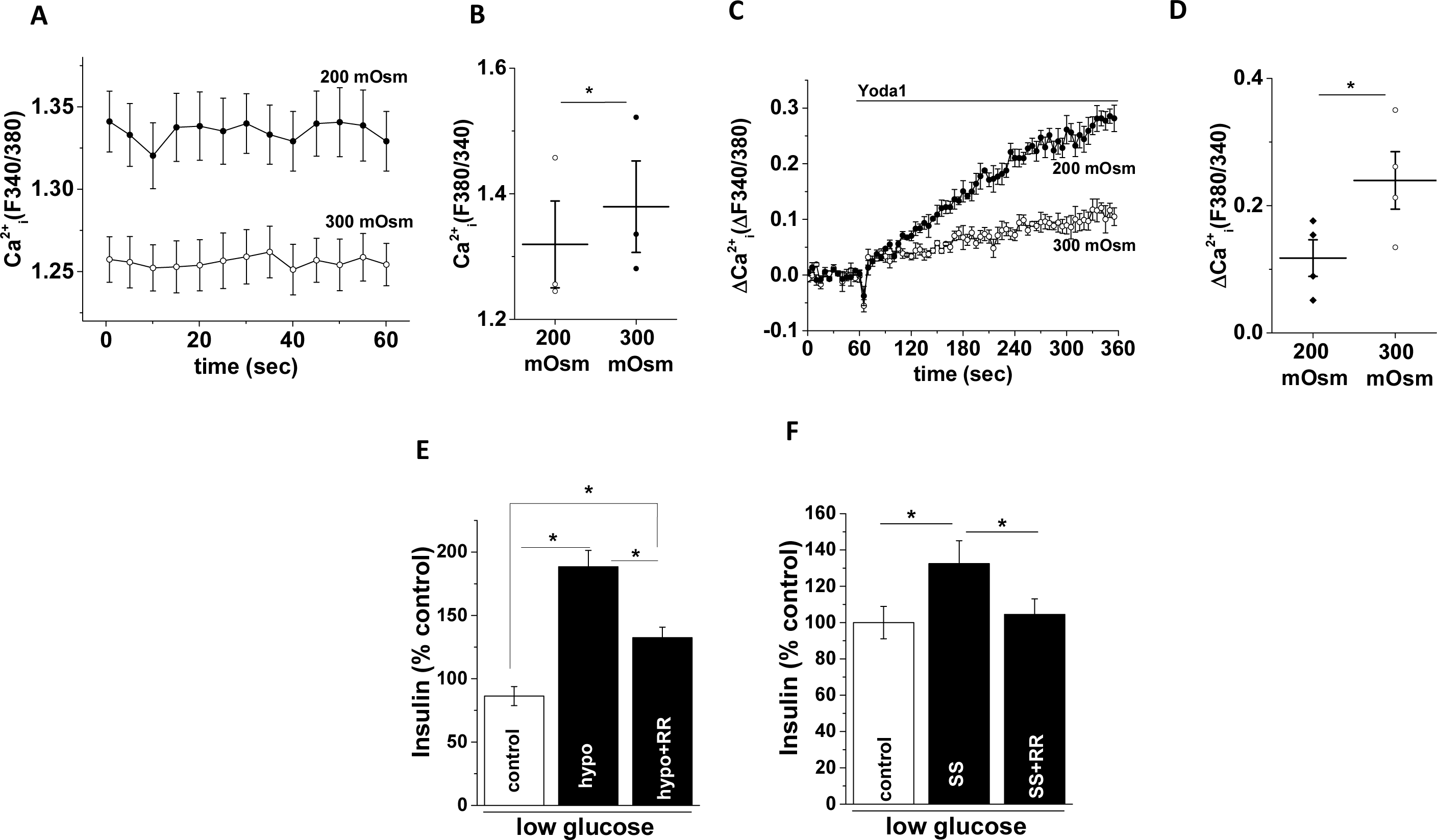
Stimulation of insulin release by osmotic or shear stress. **A)** Basal Ca^2+^ levels in INS-1 cells in hypotonic (200 mOsm) or isotonic (300 mOsm) extracellular buffer. **B)** Mean data obtained from 3 independent experiments of the types shown in (**A**). **C)** Example data showing Yoda1 or DMSO (control) induced Ca^2+^ influx in INS-1 cells in hypotonic or isotonic buffer. **D)** Mean data obtained from 3 independent experiments of the types shown in (**C**). **E)** Mean data showing insulin levels in the extracellular buffer upon treatment of INS-1 cells with hypotonic buffer for 1 hour in the presence or absence of ruthenium red (RR). **F)** As of (**E**) but showing the effect of shear stress (SS). Insulin levels were measured by ELISA, normalised against controls and expressed as a percentage (%) of control. For RR inhibition experiments, cells were pre-treated with RR for 15 min before exposure to hypotonic buffer or SS. Data are expressed as mean±sem; * denotes p<0.05; all data are representative of or mean obtained from 3 or 4 independent repeats.

### 3.4 Piezo1 agonist regulates insulin release from INS-1 cells

We next investigated the relevance of Piezo1 perturbation on high glucose induced insulin release. As expected, high glucose (17.8 mM for 1 hour) stimulated insulin release (Figure 4A). Similarly, Yoda1 (10 μM for 1 hour) stimulated insulin release in low glucose buffer (Figure 4A). The effects of high glucose and Yoda1 were not additive which may suggest either a common underlying mechanism or a limited pool of available insulin (Figure 4A). Dooku1 alone did not affect insulin release but pre-treatment with Dooku1 (10 μM for 15 min) significantly reduced the effect of Yoda1 (Figure 4A). The inactive Yoda-1 analogue (2e; 10 μM for 1 hour) did not induce insulin release. As expected, exposure to high glucose (17.8 mM for 30 min) increased basal Ca^2+^ level in INS-1 cells (Figure 4B &C). High glucose also enhanced Yoda1 induced Ca^2+^ influx (Figure 4D &E).

**Figure 4.**
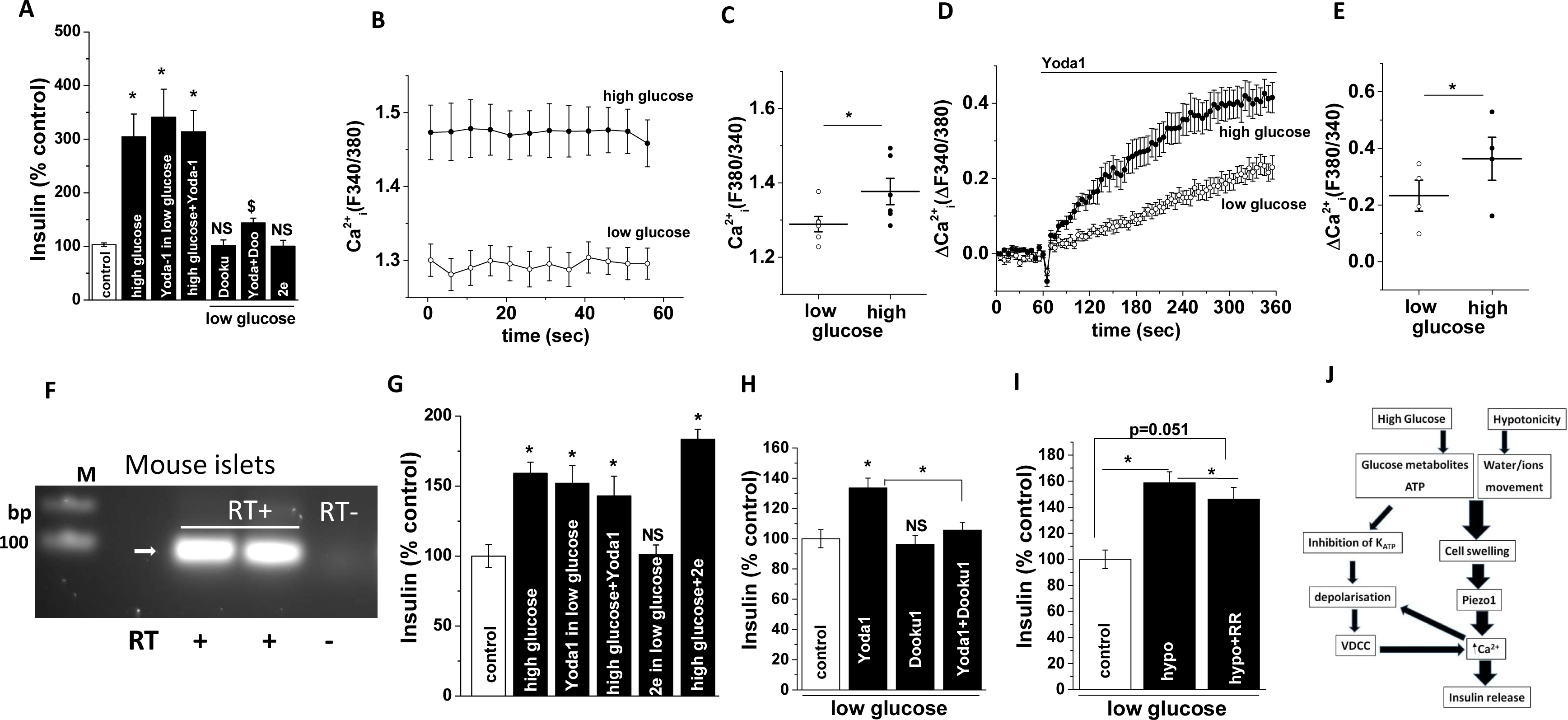
Piezo1 agonist regulates insulin release from INS-1 cells and pancreatic islets. **A)** Mean data showing insulin levels in the extracellular solution upon treatment with shown chemicals for 1 hour; 2e: non-functional Yoda1 analogue. **B)** Basal Ca^2+^ levels in INS-1 cells in low glucose (2.8 mM) or high glucose (17.8 mM) containing extracellular buffer. **C)** Mean data obtained from 3 independent experiments of the types shown in (**B**). **D)** Example data showing Yoda1 or DMSO (control) induced Ca^2+^ influx in INS-1 cells in low glucose or high glucose containing buffer. **E)** Mean data obtained from 3 independent experiments of the types shown in (**D**). **F)** Representative agarose gel electrophoresis picture with a single band at 104 bp showing Piezo1 mRNA expression (RT: with (+) and without (−) reverse transcriptase; bp: base pairs; M: marker) in mouse islets. **G)** As for (**A**) but showing data from mouse pancreatic islets. **H)** As for (**A**) but showing data from mouse pancreatic islets. **I)** Mean data showing insulin levels in the extracellular buffer upon treatment of pancreatic islets with hypotonic buffer for 1 hour in the presence or absence of ruthenium red (RR). **J)** Schematic flowchart showing possible mechanisms underlying Piezo1 mediated regulation of insulin secretion. Insulin levels were measured by ELISA, normalised against control treatments and expressed as a percentage (%) of control. For Dooku1/RR inhibition experiments, cells were pre-treated with Dooku1/RR for 15 min before the addition of Yoda1/exposure to SS. Data are expressed as mean±sem; * denotes p<0.05 vs control; NS: non-significant vs control; $ denotes p<0.05 vs Yoda1; all data are representative of or mean obtained from 3 or 4 independent repeats.

### 3.5 Piezo1 agonist regulates insulin release from primary mouse islets

To test whether Piezo1 stimulation leads to insulin release from primary β-cells we turned to isolated mouse pancreatic islets. The islets showed strong expression of Piezo1 mRNA (Figure 4F). Similar to INS-1 cells, high glucose and Yoda1 induced insulin release and the effects were not additive (Figure 4G). The inactive Yoda1 analogue (2e; 10 μM for 1 hour) had no effect (Figure 4G). When islets were pre-treated with Dooku1, the effect of Yoda1 was prevented (Figure 4H). RR inhibited hypotonicity induced insulin release from the mouse pancreatic islets as well (Figure 4I). These data suggest that Piezo1 channel activation is a mechanism for inducing insulin release.

## 4 Discussion

Ca^2+^ entry induced by Piezo1 agonist Yoda1 and the Piezo1 mRNA expression suggest the presence of functional Piezo1 channels in two types of b-cell lines – INS-1 and BRIN-BD11. This is further supported by the predicted responses to known chemical inhibitors of Piezo1 and a non-functional Yoda1 (2e) analogue in Ca^2+^ measurement experiments. Importantly, two well characterised Piezo1 activators (i.e. shear stress and Yoda1) also induced significant insulin secretion from the β-cell lines and islets. The stimulatory effects of hypotonicity, shear stress and Yoda1 were lost in the presence of Piezo1 inhibitors. Thus, the results support our hypothesis that stimulating mechanosensitive Piezo1 channels by chemical agonist or cell swelling can induce insulin secretion from pancreatic β-cells.

Yoda1 mimics mechanical stimulation and thus facilitates the study of Piezo1 channels without the need for mechanical stimulation, and has no effect on Piezo2 channels [24]. Overexpressed mouse and human Piezo1 channels were originally shown to be activated by Yoda1 with an EC_50_ of 17.1 and 26.6 μM respectively [24]. Native Piezo1 channels too have responded to Yoda1 at low micromolar concentrations [19], [21], [29]. However EC_50_ values were shown to be much lower (2.51 µM for stably overexpressed Piezo1 and 0.23 µM for native channels in human umbilical vein endothelial cells) recently [26]. In β-cells, Yoda1 induced a dose dependent increase in intracellular Ca^2+^ with an EC_50_ of 4.54 to 9 µM (Figure 1C & 2C). Furthermore, non-functional Yoda1 analogue (2e) failed to induce any Ca^2+^ influx. Both non-selective inhibitors (RR & Gd^3+^) and Dooku1 inhibited Yoda1 induced Ca^2+^ influx significantly. Despite our repeated efforts to use siRNA to silence *Piezo1* gene in β-cell lines, we were not successful in achieving efficient transfection and thus adequate Piezo1 knockdown (unpublished data). However, the concentration of Yoda1 required to stimulate Ca^2+^ entry and the lack of effect of 2e, the inhibition of Yoda1-mediated effect by removal of extracellular Ca^2+^, the expected effects of Piezo1 inhibitors and the lack of any inhibitory action of VDCC blocker (nicardipine) indicate that the Ca^2+^ influx observed in our study is predominantly through Piezo1 channels. Of note, higher concentration (>~20 μM) of Yoda1 solutions turn increasingly opaque. Therefore, the apparent EC_50_ is likely affected by compound insolubility and may not allow meaningful interpretation.

Ca^2+^ elevation upon hypotonic stimulation has already been reported in various β-cell lines and also in both primary mouse and rat pancreatic β-cells [4], [10], [13]. It was proposed that hypotonic stimulation leads to membrane depolarization because of activation of volume sensitive outwardly rectifying chloride (Cl^-^) channels which in turn activated VDCC resulting in Ca^2+^ influx and insulin release [3], [9]. In agreement, some studies have shown that hypotonically induced Ca^2+^ Ca^2+^ elevation in rat pancreatic β-cells was nearly abolished by the VDCC blocker nicardipine [4], [10]. However, a recent study in HIT clonal cells demonstrated that Cl^-^ channel blockers such as niflumic acid and DIDS failed to inhibit insulin secretion induced by hypotonic stimulation [11]. It has also been shown that hypotonically induced insulin secretion from the HC9 β-cell line is inhibited by the VDCC blockers nitrendipine and calciseptine, but not by the Cl^-^ channel blocker DIDS [6]. Notably, Gd^3+^ suppressed hypotonicity induced Ca^2+^ elevation in rat pancreatic β-cells and a role for stretch activated cation channels was proposed [13]. Hence, it is clear that there is no unanimous agreement on the exact mechanism of hypotonicity/cell swelling induced insulin release.

Having neither examined the effect of Piezo1 blockade on hypotonic response in Ca^2+^ entry nor used Cl^-^channel blockers, we can not rule either of them out. However, in the light of present data showing potentiation of Yoda1 induced Ca^2+^ entry by hypotonicity and high glucose, we propose the following model: hypotonicity induced cell swelling leading to Piezo1 activation and Ca^2+^ entry which in turn causes membrane depolarisation and VDCC activation for further Ca^2+^ entry (Figure 4J). Further careful experimentation with VDCC, Cl^-^, K_ATP_ and Piezo1 channel blockers in different combinations should be performed to clearly elucidate the existence and significance of such a pathway. However, hypotonicity induced insulin release, which could be considered as one of the read-outs for intracellular Ca^2+^ elevation is significantly, but only partially, inhibited by Piezo1 blockade (Figure 3D & 4H). Hence both pathways (chloride mediated and Piezo1/ Ca^2+^ mediated depolarisation upon cell swelling) may be participating in hypotonicity induced insulin release. Interestingly, Piezo1 channels are permeable to Cl^-^ as well [30].

Induction of β-cells to produce/release insulin using secretagogues (e.g. sulfonylureas which are K_ATP_ channel blockers) is a clinically used strategy to manage diabetes mellitus [31]. However the current secretagogue pharmaceuticals are not favoured due to their adverse cardiovascular reactions and reported induction of β-cell apoptosis [32]. The search for non-pharmacological approaches led to the identification of ultrasound as a Ca^2+^ dependent insulin secretion inducer [33], [34]. Stimulation of stretch activated channels was proposed as the underlying mechanism. Interestingly, Piezo1 can be activated by ultrasound [35]. Though the significance of Piezo1 in physiological insulin secretion is not evident from our study, using mechanosensitive Piezo1 induction as a strategy to induce insulin secretion has clear clinical potential and warrants further investigation [34]. It is an attractive approach particularly in neonatal diabetes where the disease is caused by gain-of-mutations in K_ATP_ channels [36]. However future experiments using siRNA and β-cell specific Piezo1 knockout mice are needed to clearly elucidate the role of Piezo1 in the physiological function of β-cells.

## 5 Acknowledgement

The authors thank Bhawna Chandravanshi & Prof Ramesh Bhonde (School of Regenerative Medicine, Manipal University, Bengaluru, India) for the technical assistance with mouse pancreatic islet isolation. We thank the British Heart Foundation for research grant funding.

